# Ancient DNA of *Rickettsia felis* and *Toxoplasma gondii* implicated in the death of a hunter-gatherer boy from South Africa, 2,000 years ago

**DOI:** 10.1101/2020.07.23.217141

**Authors:** Riaan F. Rifkin, Surendra Vikram, Jean-Baptiste J. Ramond, Don A. Cowan, Mattias Jakobsson, Carina M. Schlebusch, Marlize Lombard

## Abstract

The Stone Age record of South Africa provides some of the earliest evidence for the biological and cultural origins of *Homo sapiens*. While there is extensive genomic evidence for the selection of polymorphisms in response to pathogen-pressure in sub-Saharan Africa, there is insufficient evidence for ancient human-pathogen interactions in the region. Here, we analysed shotgun metagenome libraries derived from the sequencing of a Later Stone Age hunter-gatherer child who lived near Ballito Bay, South Africa, *c*. 2,000 years ago. This resulted in the identification of DNA sequence reads homologous to *Rickettsia felis*, and the reconstruction of an ancient *R. felis* genome, the causative agent of typhus-like flea-borne rickettsioses. The concurrent detection of DNA reads derived from *Toxoplasma gondii*, the causative agent of toxoplasmosis, confirms the pre-Neolithic incidence of these pathogens in southern Africa. We demonstrate that an *R. felis* and *T. gondii* co-infection, exacerbated by various additional bacterial and parasitic pathogens, contributed to the ill-health and subsequent demise of the boy from Ballito Bay.

---

The DNA of an anaemic seven-year-old boy (Pfeiffer et al., 2019), who lived in South Africa near what is today the town of Ballito Bay *c*. 2,000 years ago (ya), recently revised the genetic time-depth for *Homo sapiens* (Schlebusch et al., 2017) (Fig. 1). During the process of extracting and generating DNA data from human skeletal material, large amounts of associated genetic data are generated, *i*.*e*., shotgun meta-genome sequence data. These data contain traces of potentially pathogenic microbes associated with the person whose DNA was examined. Here, we report on the molecular detection of bacterial and parasitic pathogens associated with the boy from Ballito Bay (*i*.*e*., aDNA sample ‘BBayA’) (Supplementary Information 1 (SI)). We were able to reconstruct an ancient genome for *Rickettsia felis*, a bacterium causing typhus-like flea-borne rickettsioses. We also identified ancient DNA (aDNA) sequence reads representative of *Toxoplasma gondii*, a zoonotic protozoan intracellular parasite causing toxoplasmosis. A co-infection of these, as well as other pathogens, may well explain the boy’s reported anaemia and *cribra orbitalia* (Pfeiffer et al., 2019). *Rickettsia felis* has been widely viewed as a recent or emergent pathogen, first implicated as a cause of human illness in Texas, USA, in 1994 (Schriefer et al., 1994; Pages et al., 2010). The origin of *T. gondii* is presumed to coincide with the domestication of agricultural and companion animals following the start of the Neolithic in the Near East, *c*. 12,000 ya (Sibley, 2003). Our results show that these, and various other bacterial and parasitic pathogens, were present by at least 2,000 ya amongst southern African Stone Age hunter-gatherers who did not have domesticated animals, nor lead sedentary lives.

**Fig. 1.**
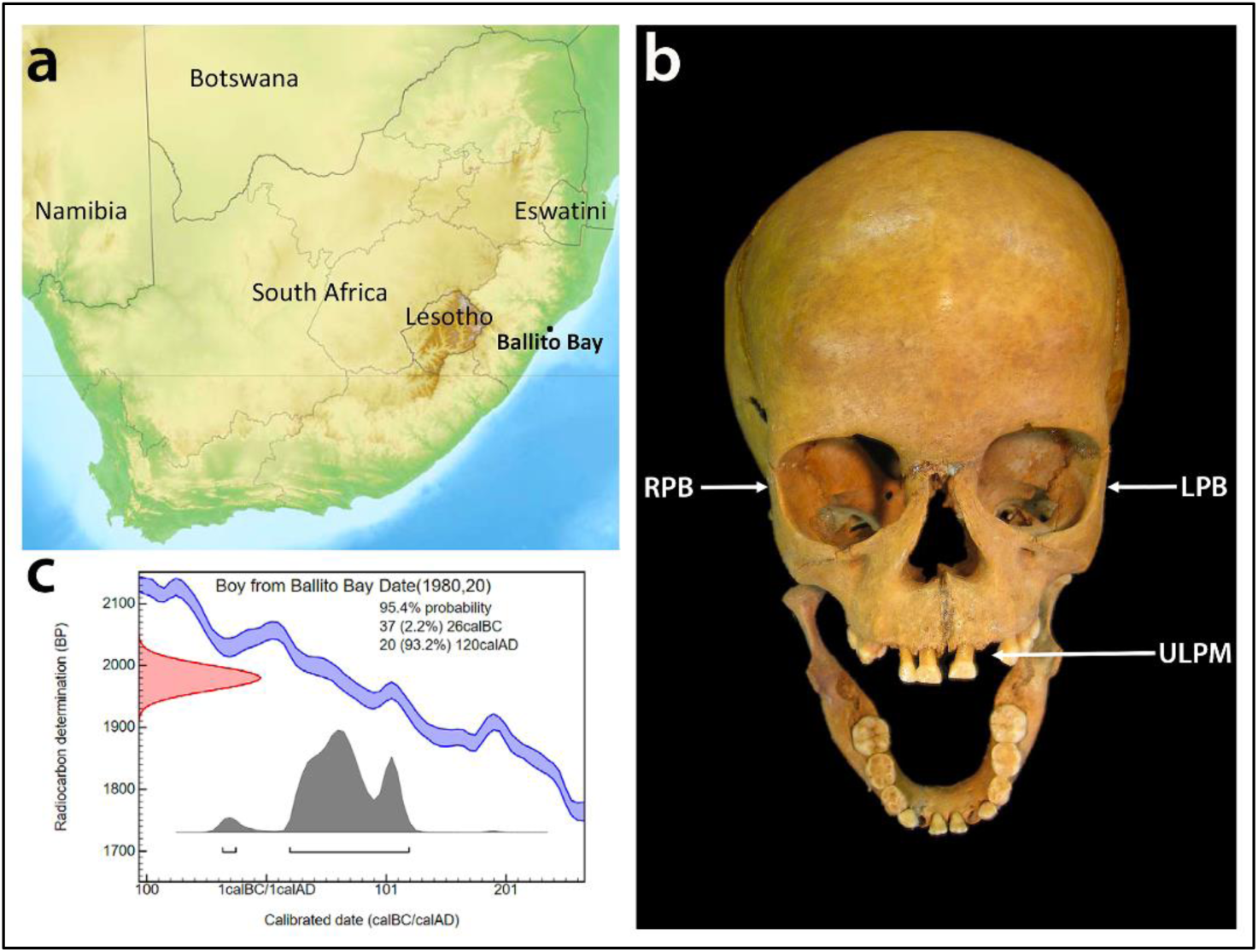
**a)** The provenience of the Later Stone Age hunter-gatherer skeletal remains recovered from a mound formed by a shell midden overlooking the beach in Ballito Bay, KwaZulu-Natal Province, South Africa. **b**) The cranial remains of the BBayA male child indicating aDNA sample sources, *i.e*., DNA was extracted and sequenced from bone samples acquired from the left petrous bone (*LPB*), right petrous bone (*RPB*) and the upper left premolar (*ULPM*). **c**) The C14 date (1,980 ± 20 cal. BP) obtained for the remains of the child.

Southern Africa has long been a hotspot for research concerning the origins of *H. sapiens* (Mounier and Lahr, 2019). The oldest genetic population divergence event of our species, at *c*. 350-260 kyr (thousand years) ago, is represented by the genome of the boy from Ballito Bay (Schlebusch et al., 2017; Lombard et al., 2018). Fossil evidence exists for early *H. sapiens* from ∼259 kyr ago (Grün et al., 1996), for late *H. sapiens* from at least 110 kyr ago (Grine et al. 2017), and for cognitive-behavioural complexity since *c*. 100 kyr ago (Henshilwood et al., 2011; Lombard, 2012; Wadley, 2015; Tylen et al., 2020). Despite the fact pathogens have long exerted a significant influence on hominin longevity (Rifkin et al., 2017) and human genetic diversity (Pittman et al., 2016), and given that diseases continue to shape our history (Andam et al., 2016), their influence on the biological and socio-cultural evolution of our species in Africa is routinely overlooked (SI 2).

The gradual dispersal of *H. sapiens* from Africa into Asia and Europe was accompanied by various commensal and pathogenic microbes (Houldcroft et al., 2017; Reyes-Centeno et al., 2017; Pimenoff et al., 2018). The presence of specific TLR4 polymorphisms (*i*.*e*., pathogen-recognition receptors) in African, as well as in Basque and Indo-European populations, suggests that some mutations arose in Africa before the dispersal of *H. sapiens* to Eurasia (Ferwerda et al., 2008). In addition, the bio-geographic distribution of *Plasmodium falciparum* (Tanabe et al., 2010) and *Helicobacter pylori* (Linz et al., 2007) exhibits declining genetic diversity, with increasing distance from Africa, with ‘Out of Africa’ estimates of ∼58 kyr and ∼80 kyr ago, respectively. From records such as these, it is apparent that persistent exposure to pathogens exerted selective pressure on human immune-related genes (Nédélec et al., 2016; Owers et al., 2017; Schlebusch et al., 2020), cognitive development (Kessler et al., 2017) and social behaviour (Thornhill and Fincher, 2014). The adverse influence of diseases on ancient forager populations is exemplified by the fact that infectious, zoonotic and parasitic diseases account for ∼70% of deaths recorded amongst contemporary hunter-gatherers (Gurven and Kaplan, 2007) (SI 2).

While there is substantial evidence for the selection of human genomic polymorphisms in response to pathogen-pressure in sub-Saharan Africa (SI 2), there is little direct evidence of ancient human-pathogen interactions in the region. To gain insight into the prehistoric incidence of human pathogens, we analysed eight shotgun meta-genome libraries originating from the sequencing of the boy from Ballito Bay (Fig. 1). We were able to identify, at species level, 515,174 unique authenticated aDNA sequence reads mapping to the genome of *R. felis* strain LSU-Lb (SI 3), and 769,584 unique reads partially mapping to the *T. gondii* TgCatPRC2 reference genome (SI 4) (Table 1). At the genus level, we identified aDNA reads homologous to *Anaplasma, Babesia, Bordetella, Brucella, Leishmania, Plasmodium* and *Trypanosoma* (Table S1). Taxonomic classification of metagenomic reads was achieved using Kraken2 (Wood et al., 2019) and a custom database of bacterial, archaeal, protozoal and viral genomes from the NCBI RefSeq database (https://www.ncbi.nlm.nih.gov/refseq/).

**Table 1.** Authenticated ancient DNA sequence reads, derived from eight aDNA shotgun metagenome sequence libraries generated from the boy from Ballito Bay (BBayA), mapped to the *Rickettsia felis* LSU-Lb and *Toxoplasma gondii* TgCatPRC2 reference genomes.

|  | Sample source | Libraries | Total reads | Human reads | *Rickettsia* reads | % of reads | % duplicates | Unique *R. felis* reads | Mean read length (bp) | Genome coverage (x) |
| --- | --- | --- | --- | --- | --- | --- | --- | --- | --- | --- |
| *Rickettsia felis* | Left petrous bone | 5 | 3768601170 | 491223634 | 525855 | 0.0140 | 37.53 | 328461 | 84.40 | 18.4353 |
|  | Right petrous bone | 2 | 1075114194 | 244053412 | 250195 | 0.0233 | 25.57 | 186219 | 84.90 | 10.5154 |
|  | Upper left premolar | 1 | 42434936 | 18050 | 562 | 0.0013 | 12.09 | 494 | 83.00 | 0.0001 |
|  | **Total** | **8** | **4886150300** | **735295096** | **776612** | **0.0159** | **33.66** | **515174** | **84.57** | **28.9735** |
|  | Sample source | Libraries | Total reads | Human reads | *Toxoplasma* reads | % of reads | % duplicates | Unique *T. gondii* reads | Mean read length (bp) | Genome coverage (x) |
| *Toxoplasma gondii* | Left petrous bone | 5 | 3768601170 | 491223634 | 9811214 | 0.2603 | 95.68 | 423449 | 96.74 | 0.5016 |
|  | Right petrous bone | 2 | 1075114194 | 244053412 | 4288620 | 0.3989 | 91.94 | 345635 | 90.85 | 0.3851 |
|  | Upper left premolar | 1 | 42434936 | 18050 | 521 | 0.0012 | 4.03 | 500 | 69.44 | 0.0004 |
|  | **Total** | **8** | **4886150300** | **735295096** | **14100355** | **0.2886** | **94.54** | **769584** | **94.08** | **0.8871** |

Pathogenic taxa were identified, and their reference genomes downloaded from the NCBI RefSeq database for downstream analysis. The mapping of candidate taxa was performed on a competitive basis against bacterial and parasitic genomes, and a complete human genome, *i*.*e*., *H. sapiens* assembly version GRCh38/hg39 (Table S2) (Materials and Methods). The authentication of aDNA reads ascribed to these taxa was achieved by library-independent verification using mapDamage (Jónsson et al., 2013) and the analyses of the read-length distribution (bp) (Fig. S1).

To confirm that the organism represented in our metagenomic output was an *R. felis* strain, and not one of its closely related southern African species (*e*.*g*., *R. prowazekii, R. typhi, R. conorii* and *R. africae*), and to detect signs of plasmid rearrangements, we mapped our datasets against all currently available (*i*.*e*., 126) NCBI *R. felis* reference genomes (Table S3). The plasmid system in *R. felis* is unusual since no other bacteria in the Rickettsiales (*i*.*e*., *Anaplasma, Neorickettsia* and *Wolbachia*) are known to harbour plasmids. *Rickettsia* has small genomes comprising 1.1-1.8 million base pairs (Mbp) and a high percentage of non-coding DNA, indicative of a process of reductive evolution (Gillespie et al., 2015).

Their unique genomic structure nevertheless allows high mapping specificity across the *R. felis* genome, which in turn, allowed us to infer the presence and absence of genomic regions from the level of coverage observed after mapping the raw datasets to the reference genomes (Fig. 2a). We were able to assemble ∼99.90% of the *R. felis* chromosome at a mean depth of coverage of 11.41-fold. The BBayA dataset contains an *R. felis* strain remarkably similar to both the *R. felis* URRWxCal2 (GCA_000012145.1) and *R. felis* LSU-Lb (GCA_000804505.1) reference genomes, with an average nucleotide identity (ANI) of 99.95% and 99.90%, respectively (Table S4). Subsequent phylogenetic analysis revealed that the *R. felis* LSU-Lb strain is the closest homologue to the ancient BBayA *R. felis* strain described here.

**Fig. 2.**
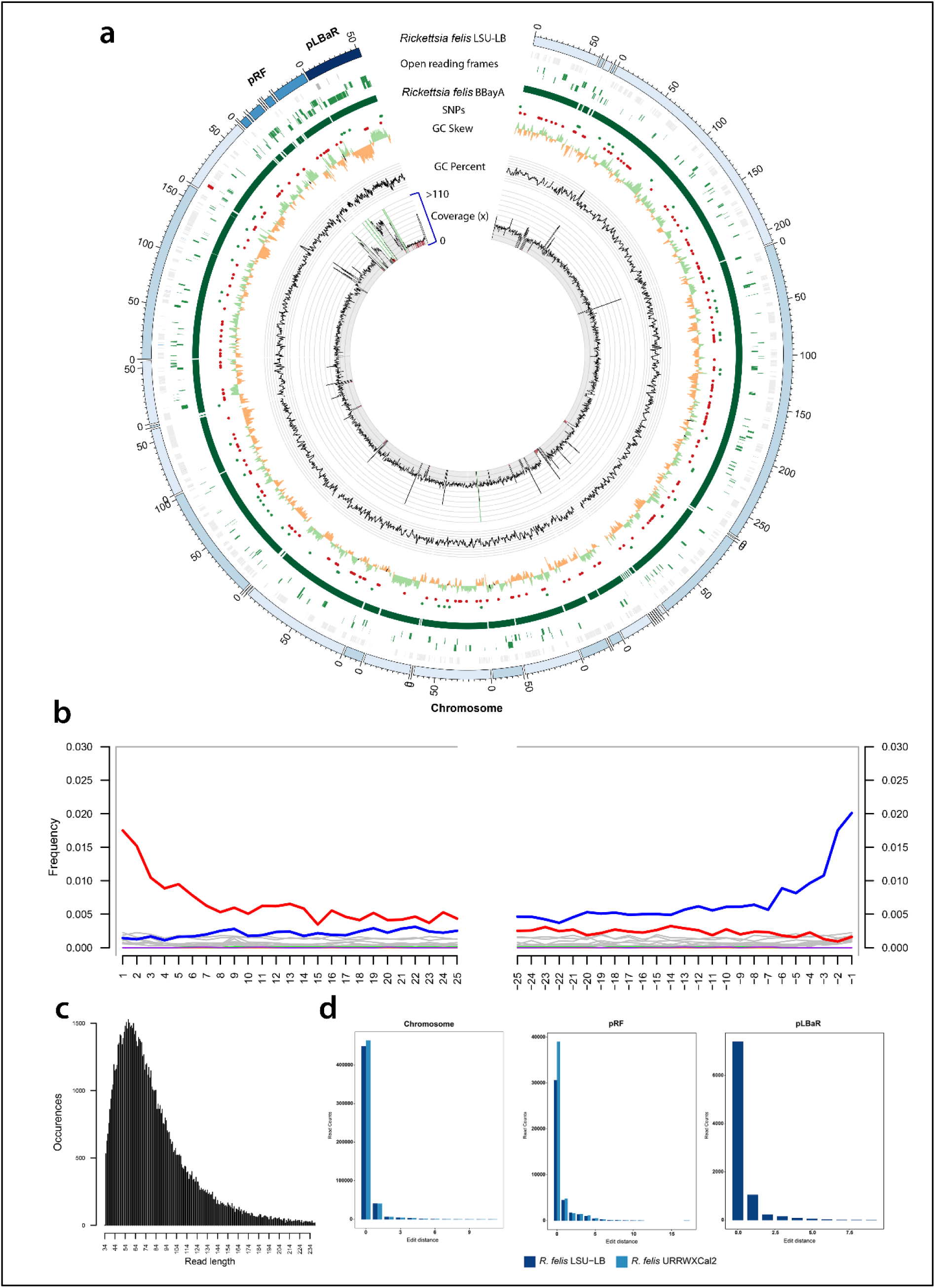
**a)** Genome reconstruction of the ancient BBayA *R. felis* genome and mapping of the ancient genome to the genome of *R. felis* LSU-Lb. The comparison was performed with the chromosome and plasmids (*pRF* and *pLBaR*). Rings (from outer to inner ring) show ORFs, SNPs, GC skew, GC content and coverage. Green and red dots represent SNPs on the positive or negative DNA strands, respectively. Following the initial mapping of our datasets against all available (*i.e*., 126) NCBI *R. felis* reference genomes (Table S3), the genome coverage analysis of BBayA *R. felis* was performed using the reads mapping to the *R. felis* LSU-Lb genome (the closest phylogenetic homologue to the ancient BBayA *R. felis* strain), with average coverage estimated at 28.97-fold. **b**) DNA damage pattern analysis for the BBayA *R. felis* using mapDamage. G-to-A and C-to-T misincorporations are plotted in blue and red, respectively, and grey lines indicate all possible misincorporations. **c**) DNA fragment read-length distributions of the BBayA *R. felis* reads, exhibiting a mean read-length of 82.30 base-pairs (bp). **d**) Distribution of edit distance of high quality BBayA *R. felis* reads mapping to *R. felis* LSU-Lb and *R. felis* URRWxCal2. The plasmid *pLbaR* occurs only in *R. felis* LSU-Lb and BBayA, with *R. felis* URRWxCal2 devoid of this plasmid.

The reconstruction of the BBayA *R. felis* genome resulted in the recognition of the single *Rickettsia* chromosome and the detection of two plasmids, *i*.*e*., *pRF* and *pLbaR* (Table S4). The authentication of aDNA sequence reads ascribed to *R. felis* was achieved by library-independent verification using mapDamage (Jónsson et al., 2013) (Fig. 2b) and analyses of the read-length distribution (bp) (Fig. 2c) and edit-distances (Fig. 2d). Consistent with aDNA, we detected significant DNA damage patterns for the reads mapping to the *R. felis* genome assembly (SI 5). The mean read-length distribution of all BBayA *R. felis* datasets (84.57 bp) furthermore indicated that the DNA was in a highly fragmented state. Damage pattern and read-length distribution analysis of the host (BBayA) DNA exhibited a similar DNA damage profile and short (*i*.*e*., damaged) sequence read-length distribution (Fig. S2).

Because *R. felis* displays genotypic and phenotypic attributes of both spotted fever (SFG) and typhus group (TG) rickettsiae, it is difficult to place phylogenetically (Fig. S3). Phylogenetic analyses of the BBayA *R. felis* genome revealed clear clustering within the recently classified *R. felis* transitional group *Rickettsia* (TRG), which is characterised by including both vertebrate *Rickettsia* and *Rickettsia* infecting non-blood feeding arthropods (Fig. 3). Specifically, the ancient BBayA *R. felis* genome occurs between *R. felis* LSU and *R. felis* LSU-Lb within the transitional group (TRG) rickettsiae, with close phylogenetic affinities to the reference *R. felis* URRWxCal2 reference genome. BBayA *R. felis* also exhibits close affinities to the better-known and highly pathogenic *R. typhi* (the causative agent of murine typhus) and *R. prowazekii* (the etiologic agent of epidemic typhus) (Fig. 2).

**Fig. 3.**
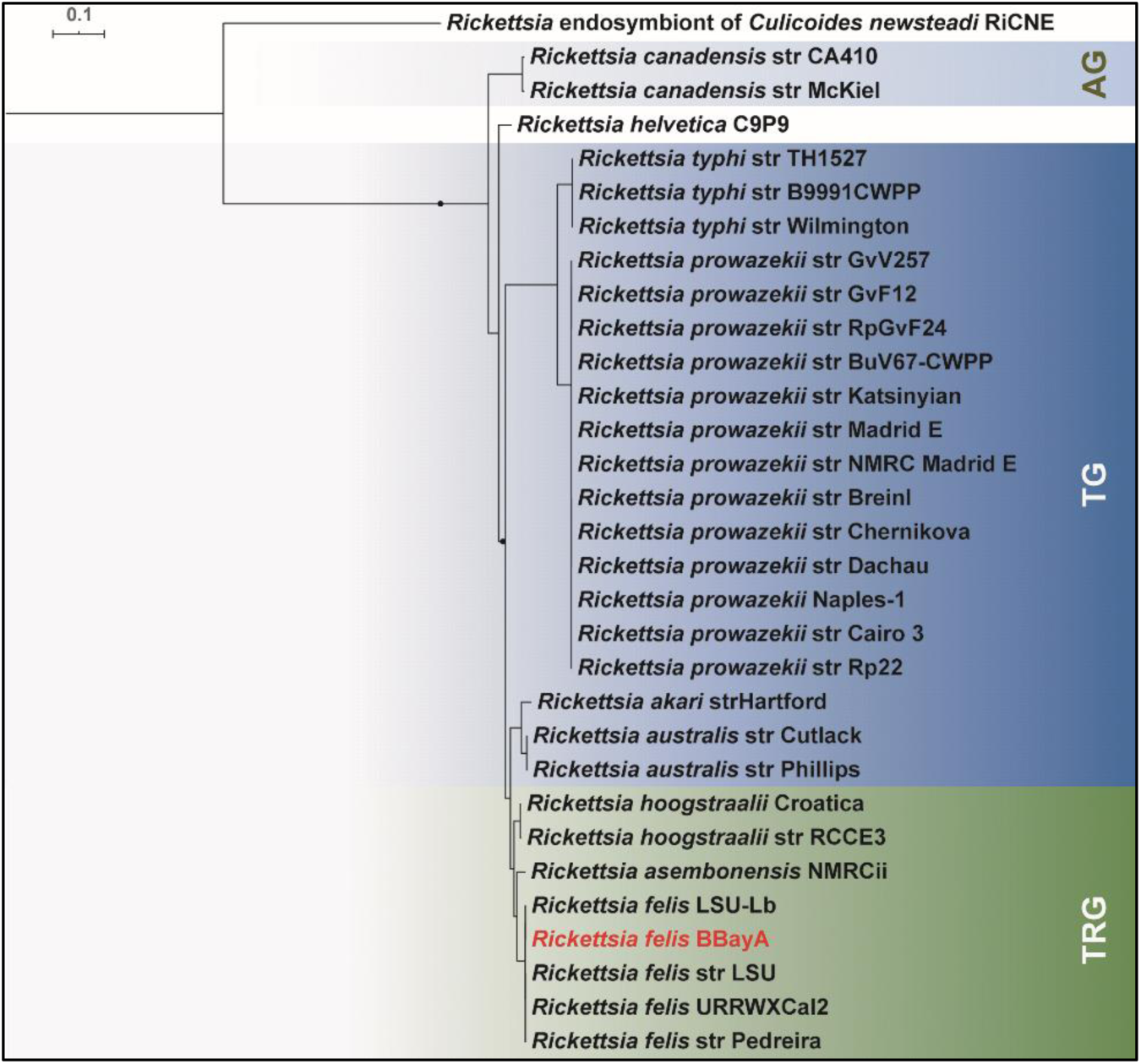
Representation of the phylogeny of the ancient *R. felis* strain derived from BBayA (*Rickettsia felis* BBayA) indicated within the context of the *Rickettsia* ‘typhus’ (TG) and ‘transitional’ (TRG) groups. Single nucleotide polymorphisms (SNPs) were combined with all other SNPs found in *R. felis* strains included in this study to construct a phylogenetic tree using the maximum likelihood method. The legend displays the branch length. The phylogenetic tree was constructed using the best model predicted by JmodelTest (Posada, 2008). The maximum-likelihood tree was prepared using the GTR+G+I substitution model and visualised in iTOL tree-viewer (Letunic and Bork, 2007).

The chronometric date of ∼2,000 years for the skeletal remains enabled us to date stages in the evolution of the BBayA *R. felis* using a Bayesian and Markov Chain Monte Carlo phylogenetic approach (Materials and Methods). Our analyses indicated a divergence estimate of *c*. 2.5 (2.63 −2.31) million years ago (mya) for *R. felis* from the globally distributed and particularly pathogenic *R. prowazekii* and *R. typhi* (Fig. S3). This period is marked by the emergence of the first member of our genus, *H. habilis* (Spoor et al., 2015), the very first evidence for the consumption of meat and marrow from large fauna (Domínguez-Rodrigo et al., 2010) and the initial dispersal of hominins from Africa (Scardia et al., 2019). Our analyses also revealed the emergence of a most recent common ancestor (MRCA) for the southern African *R. felis* group at 5,000 ya (*i*.*e*., 5,282 ya - 4704 ya), during the microlithic Wilton techno-complex of the southern African Later Stone Age (LSA) (Lombard et al., 2012) (SI 6). The divergence time for *R. felis* URRWxCal2 and *R. felis* BBayA was estimated at 2,942 ya and that of *R. felis* LSU-Lb and *R. felis* BBayA at *c*. 2,000 ya (Fig. S3) (Materials and Methods).

When compared to the *R. felis* (*i*.*e*., LSU-Lb and URRWxCal2) and other *Rickettsia* genomes used in this study (*i*.*e*., *R. typhi, R. prowazekii* and *R. africae*), several SNPs are specific to the BBayA *R. felis* strain (Table S4). One missense variant (mutation) was identified in the *cell surface protein 2* (Sca2) coding region of *R. felis* LSU-Lb, but was absent in URRxCal2. Sca2 (*pRF*25) was detected on the BBayA *R. felis pRF* plasmid. It is a noteworthy virulence protein in *Rickettsia* as it facilitates cell adherence (Cardwell and Martinez, 2009) and promotes pathogenesis in primary and secondary hosts (Gillespie et al., 2015). In addition, the plasmid *pLbaR* encodes a repeats-in-toxin-like type I secretion system and an associated RHS-like toxin, namely *pLbaR-*38.

As with *R. felis*, the authentication of unique DNA sequence reads ascribed to *T. gondii* (144,771) was achieved by library-independent verification using mapDamage (Jónsson et al., 2013) and analyses of the read-length distribution (bp) (Fig. S4). The mean read-length distribution of the BBayA *T. gondii* datasets (94.08 bp) showed that the DNA was in a highly fragmented state, consistent with what is expected when working with aDNA. Since our ancient *T. gondii* sequence reads mapped largely to repeat regions in the reference genome (TgCatPRC2) and not to coding gene regions (with average coverage estimated at only 0.88-fold), in-depth genomic and phylogenetic analyses of the BBayA *T. gondii* could not be performed.

Our results indicate that an *R. felis* and *T. gondii* co-infection (SI 7), exacerbated by members of at least seven additional bacterial (*i*.*e*., *Anaplasma, Bordetella* and *Brucella*) and parasitic (*Babesia, Leishmania, Plasmodium* and *Trypanosoma*) genera, contributed to the frailty and death of the boy from Ballito Bay (SI 8). Although discerning between closely related species with similar genomic elements is challenging, taxonomic classification using a high confidence value (*i*.*e*., 0.85) and species-similarity value (*i*.*e*., *k*=35) suggests that these genera are, in all probability, represented by at least some pathogenic species (Table S2) (Materials and Methods). Granting that the molecular- and population-level interactions between multiple infecting pathogenic microbes are variable and complex (Hamelin et al., 2019), the scenario of a co-infection of the child would certainly have exacerbated the clinical progression of his infection, thus accelerating his morbidity and mortality (Griffiths et al., 2011). Given reported disease case fatality rates (CFRs) for the pathogenic microbes identified in the BBayA child, the severity of his co-infective clinical condition becomes apparent (Table S5) (SI 9).

## Implications of the molecular detection of various 2,000-year-old pathogens in association with a South African hunter-gatherer child

Formerly, the identification of skeletal pathologies used to be the only means by which information concerning ancient diseases could be gained. The increasingly successful extraction of DNA from ancient human remains, and the available sequencing and bioinformatic data-processing technologies, is rapidly advancing our understanding of the antiquity of human-pathogen interactions. It has been demonstrated that the DNA of pathogenic bacteria, such as *Brucella melitensis* (Kay et al., 2014), *Mycobacterium leprae* (Schuenemann et al., 2013), *M. tuberculosis* (Müller et al., 2014), *Yersinia pestis* (Rasmussen et al., 2015), *Salmonella enterica* (Vågene et al., 2018) and *Borrelia recurrentis* (Guellil et al., 2018), viruses, such as Hepatitis B virus (Patterson Ross et al., 2015), and parasitic organisms including *Plasmodium falciparum* (Marciniak et al., 2016), can be retrieved from ancient human skeletal remains. Here, we add *R. felis, T. gondii* and several additional taxa, including members of the genera *Anaplasma, Babesia, Bordetella, Brucella, Leishmania* and *Trypanosoma* to the list of pathogens than can be recovered from ancient African human remains.

Our results provide insight into the most plausible range of pathogenic microbes implicated in the death of the boy from Ballito Bay (SI 9). Osteobiographic analysis (Pfeiffer et al., 2019) is consistent with the premise that various chronic and acute viral, bacterial and parasitic infections could have produced the skeletal signs of anaemia observed in the child (Jonker et al., 2017). Indications of *cribra orbitalia* are a symptom of marrow expansion caused by haemopoietic factors, and has been attributed to both malnutrition (*e*.*g*., megaloblastic anaemia) and parasitism (Pfeiffer et al., 2019).

Other plausible causes for this pathology include malaria (*Plasmodium sp*.), hookworm infection (*Ancylostoma duodenale* and *Necator americanus*) and schistosomiasis (*Schistosoma haematobium*), the latter of which was suggested as the best-fit cause for the child’s pathology (Pfeiffer et al., 2019). In addition to *R. felis* and *T. gondii*, both of which cause comparable osteological pathologies (Briggs et al., 2016; Weiss and Dubey, 2009), and although we did recover genetic traces of *Plasmodium*, the parasitic pathogens referred to above (Pfeiffer et al., 2019) were absent from our dataset. Instead, we recovered evidence for co-infection with members of the genera *Anaplasma, Babesia, Bordetella, Brucella, Leishmania* and *Trypanosoma* (SI 8). These, and also *R. felis* and *T. gondii*, are obligate intracellular pathogens, all of which modifies the cytoskeletal architecture and the endomembrane system of their host cells to establish productive infections (Romano and Coppens, 2013).

Here, we have demonstrated that the BBayA *R. felis* MRCA pre-dates 5,000 ya, and that the strain that infected the boy from Ballito Bay diverged locally from other TRG *R. felis* strains ∼3,000 ya. *Rickettsia felis* can, therefore, no longer be considered a novel or emerging pathogen that originated in the global north. There also appears to have been no major changes in either the virulence or host specificity, over the past ∼2,000 years of evolutionary history of *R. felis* in southern Africa. Since the sequence data suggest that the genomes of TRG *R. felis* remained largely stable over at least three millennia, having diverged only 0.05% and 0.10 % from the *R. felis* URRWxCal2 and *R. felis* LSU-Lb reference genomes, the variation that exists between extant genomes may represent transient genetic fluctuation, the evolutionary relevance of which is still uncertain (Zhou et al., 2018). Conversely, the typically small genomes of *Rickettsia* (1.1-1.8 million bp) and a high percentage of non-coding DNA (Gillespie et al., 2015) may also explain the limited divergence observed. Although tentative, our results also suggest that *R. felis* does not appear to have evolved to become either more human-adapted, or more virulent, as is the case with *Salmonella enterica* and which is associated with the cultural and economic transformations following the beginning of the Neolithic (Key et al., 2020).

With regards to the pathogenicity of the BBayA *R. felis* strain, the presence of both the Sca2 (*pRF*25) and the RHS-like toxin (*pLbaR*-38) mutations suggests that this ancient strain was, in all probability, just as pathogenic as current *R. felis* variants, and that it may well have resulted in symptoms typical of typhus-like flea-borne rickettsioses, including fever, fatigue, headache, maculopapular rash, sub-acute meningitis and pneumonia. In Africa, *R. felis* is the causative organism of many (∼15%) cases of illnesses classified as ‘fevers of unknown origin’, including febrile seizures or convulsions (Briggs et al., 2016). Relative to TG (*i*.*e*., transmitted by body lice and fleas) and SFG (transmitted by ticks) rickettsiae, a much wider host range has been reported for TRG rickettsiae, including ticks, mites, fleas, booklice and various other haematophagous insects (Gillespie et al. 2015), including mosquitos of the genera *Aedes* and *Anopheles* (Parola et al., 2016). In addition, similar to *R. typhi, R. felis* is also shed in flea faeces, providing an additional avenue for zoonotic host to human infection.

Although the emergence of *T. gondii* as a human parasite is generally associated with animal domestication (Sibley, 2003), our study demonstrates that *T. gondii* was present in a child whom followed a forager lifeway before the arrival, in South Africa, of pastoralists originating from east Africa, and farmers from West Africa (Schlebusch et al., 2017; Schlebusch and Jakobsson, 2018). In this context, human *T. gondii* infection does not therefore relate to the expansion of agriculture and its association with domestic cats (*Felis catus*) and house mice (*Mus musculus*) (Shwab et al., 2018).

Instead, the transmission of *T. gondii* to humans may originally have occurred via wild felids, the only definitive hosts of *T. gondii*, through what is termed the sylvatic life cycle (Shwab et al., 2018). The geographic distribution of African *T. gondii* genotypes indicate a separation of unique and non-classical genotypes (*i*.*e*., *Africa 1* and *Africa 3*) between the arid zones of North-East Africa and the tropical zone in West-Central Africa, with the latter region conceivably representing a refuge zone for as yet unknown and, possibly, ancestral *T. gondii* strains (Galal et al. 2019). Since early African hominins (*e*.*g*., *Australopithecus africanus, H. habilis* and *H. erectus*), and also later behaviourally-modern hunter-gatherers (*H. sapiens*) competed with indigenous African felines (*e*.*g*., *Panthera* sp.) as apex predators for at least 2 to 3 mya (Faurby et al., 2020), it is possible that both the human host-specificity and the behaviourally-manipulative abilities of *T. gondii* (Poirotte et al. 2016) evolved in the human lineage long before animal domestication (SI 7).

Whereas the first description of typhus-like disease appears in AD 1489, during the War of Granada (Pages et al., 2010), there is no previously reported evidence for the presence of *T. gondii* in either antiquity, or in prehistory. Our findings provide novel baseline data on the incidence of various pathogenic microbes amongst ancient, pre-Neolithic, southern African hunter-gatherers, necessitating further discussion about the susceptibility to, and the population impacts of, zoonotic diseases on human longevity and behaviour in the past. Globally, hunter-gatherer social networks have been shown to facilitate both the transmission and the persistence of various infectious, zoonotic and parasitic diseases (Henn et al., 2012), therefore preventing a reduction in infection risk which is generally expected to have occurred amongst presumably isolated and itinerant hunter-gatherers (Gurven and Kaplan, 2007; Houldcroft et al., 2017). It is evident that, given the temporal depth of human occupation in sub-Saharan Africa, and the preservation of DNA in local archaeological contexts, the region is well positioned to play a key role in the exploration of ancient pathogenic drivers of human evolution and mortality.

## Acknowledgements

RFR acknowledges funding provided by a National Geographic Society Scientific Exploration Grant (No. NGS-371R-18) and by the Oppenheimer Endowed Fellowship in Molecular Archaeology (the Benjamin R. Oppenheimer Trust). CMS is funded by the European Research Council (ERC) under the European Union’s Horizon 2020 Research and Innovation Programme (Grant Agreement No. 759933) and the Knut and Alice Wallenberg Foundation. We thank Yves Van de Peer and Stephane Rombauts (Bioinformatics and Evolutionary Genomics Group, VIB-UGent, Ghent, Belgium) and Ansie Yssel (BGM, CMEG, University of Pretoria, Pretoria, South Africa) for informative discussions and analytical support. Sequencing was performed at the SNP&SEQ Technology Platform, SciLife Lab, National Genomics Infrastructure, Uppsala, and computational analyses were performed at the Centre for Microbial Ecology and Genomics (CMEG), University of Pretoria, South Africa.

## Author contributions

RFR, ML and JBR conceived the study and composed the manuscript. SV and RFR performed the bioinformatic and statistical analyses, and RFR, ML and SV generated the figures. CS, MJ and ML generated the sequence datasets and DAC provided access to analytical fascilities. The KwaZulu-Natal museum provided access to the human remains in terms of sampling, export and dating permits issued to ML (#s 0014/06, 1939, 1940) according to the KwaZulu-Natal Heritage Act No. 4 of 2008 and Section 38 (1) of the National Heritage Resources Act No. 25 of 1999. Final reports have been submitted to the repository and both heritage agencies. All authors contributed to the completion of the final manuscript. All authors read and approved the final manuscript.

## Materials and Methods

### aDNA sources and extraction

Information concerning the sampling protocol is provided in Schlebusch et al. (2017). Briefly, the bone samples were UV irradiated (254 nm) for 30 minutes to one hour per side and stored in plastic zip-lock bags until sampled. Further handling of the specimens was done in a bleach-decontaminated (DNA Away, ThermoScientific) enclosed sampling tent with adherent gloves (Captair Pyramide portable isolation enclosure, Erlab). Teeth were wiped with 0.5% bleach (NaOH) and UV-irradiated sterile water (HPLC grade, Sigma-Aldrich). The outer surface was removed by drilling at low speed using a portable Dremel 8100, and between 60 and 200 mg of bone powder was sampled for DNA analyses from the interior of the bones and teeth. The researchers wore full-zip suits with caps, facemasks with visors and double latex gloves and the tent was frequently cleaned with DNA-away during sampling.

The 1.5 ml tubes containing the bone powder samples were thoroughly wiped with DNA-away before they were taken into the dedicated aDNA clean room facility at Uppsala University (Schlebusch et al., 2017). The laboratory is equipped with, among other things, an air-lock between the lab and corridor, positive air pressure, UV lamps in the ceiling (254nm) and HEPA-filtered laminar flow hoods. The laboratory is frequently cleaned with bleach (NaOH) and UV-irradiation and all equipment and non-biological reagents are regularly decontaminated with bleach and/or DNA-away (ThermoScientific) and UV irradiation. DNA was extracted from between 60 and 190 mg of bone powder using silica-based protocols, either as in Yang et al. (1998) with modifications as in Malmström et al. (2007) or as in Dabney et al. (2013), and were eluted in 50-110 μl Elution Buffer (Qiagen). Between 3 and 6 DNA extracts were made for each individual (or accession number) and one negative extraction control was processed for every 4 to 7 samples extracted.

The optimal number of PCR cycles to use for each library was determined using quantitative PCR (qPCR) in order to see at what cycle a library reached the plateau (where it is saturated) and then deducting three cycles from that value. The 25 µl qPCR reactions were set up in duplicates and contained 1 µl of DNA library, 1X Maxima SYBR Green Mastermix and 200 nM of each IS7 and IS8 primers (Meyer and Kircher, 2010) and were amplified according to supplier instructions (ThermoFisher Scientific). Each library was then amplified in four or eight reactions using between 12 and 21 PCR cycles. One negative PCR control was set up for every four reactions. Blunt-end reactions were prepared and amplified as in Günther et al. (2015) using IS4 and index primers from Meyer and Kircher (2010). Damage-repair reactions had a final volume of 25 μl and contained 4 μl DNA library and the following in final concentrations; 1X AccuPrime Pfx Reaction Mix, 1.25U AccuPrime DNA Polymerase (ThermoFisher Scientific) and 400nM of each the IS4 primer and index primer (Meyer and Kircher, 2010). Thermal cycling conditions were as recommended by ThermoFisher with an annealing temperature of 60°C (Meyer and Kircher, 2010). The negative controls processed did not yield any DNA and were therefore not sequenced (Schlebusch et al., 2017). The DNA libraries were sequenced at SciLife Sequencing Centre in Uppsala using either Illumina HiSeq 2500 with v2 paired-end 125 bp chemistry or HiSeq XTen with paired-end 150 bp chemistry.

### Authentication of ancient pathogenic DNA

Following the application of bioinformatic analytical protocols, the resultant data-set indicated the presence of a single authentic (ancient) pathogenic taxon subjected to and verified according to the authentication process outlined above in Schlebusch et al. (2017). Briefly, molecular damage accumulating after death is a standard feature of all aDNA molecules. The accumulation of deaminated cytosine (uracil) within the overhanging ends of aDNA templates typically results in increasing cytosine (C) to thymine (T) misincorporation rates toward read starts, with matching guanine (G) to adenine (A) misincorporation rates increasing toward read ends in double-stranded library preparations (Briggs et al., 2007). Being the ‘gold-standard’ of aDNA authentication, we used mapDamage v2.0.1 (Jónsson et al., 2013) to determine the incidence of cytosine (C) to thymine (T) and guanine (G) to adenine (A) substitution rates at the 5’-ends and 3’-ends of strands (Briggs et al., 2007).

### Sequence data processing and analysis

Paired-end aDNA sequencing reads were first processed to facilitate the removal of adapters and primers using AdapterRemoval v2 (Schubert et al., 2016) following the parameters ‘min-quality’ 20, ‘min-length’ 35 and ‘collapsed to merge’ the forward- and reverse-sequence reads. Human (*i*.*e*., *H. sapiens*) reads were removed using the BWA-MEM algorithm against the human reference genome (Li and Durbin, 2010). Kraken2 analysis (Wood et al., 2019) was performed using a custom database (including selected bacterial, archaeal, protozoal and viral taxa) derived from the NCBI RefSeq database (https://www.ncbi.nlm.nih.gov/refseq/) with a high confidence (*i.e*., ‘cut-off’ level) value of 0.85 to obtain the most accurate taxonomic assignments. The identification of microbial taxa is based on the use of exact-match database queries of *k*-mers, instead of alignment similarity. As different ‘*k*’ values approximate degrees of taxonomic similarity, with *k*=21 indicative of genus-level similarity, *k*=31 of species-level similarity and *k*=51 of strain-level similarity, we applied the default *k* value setting of 35 (*i.e*., *k*=35). Using these results, pathogenic taxa were identified, and their respective reference genomes downloaded from the NCBI RefSeq database for the downstream analysis. Competitive alignment with BWA v0.6.2-r126 (Langmead and Salzberg, 2012) was performed using the eight BBayA aDNA sequencing libraries (*i.e*., the ‘petrous left’, ‘petrous right’ and ‘premolar’ DNA sample libraries). Exact duplicates were removed using MarkDuplicates (Picard) (https://gatk.broadinstitute.org/hc/en-us/articles/360037052812-MarkDuplicates-Picard-).

### Genome reconstruction and comparative analysis of BBayA *R. felis*

The *R. felis* LSU-LB and URRWXcal2 strains were used as reference genomes during the BWA v0.6.2-r126 alignment to the BBayA *R. felis* chromosomes and plasmids. FASTQ reads were extracted from the resulting alignment and *de-novo* assembly was performed using the SPAdes v3.11 genome assembler (Bankevich et al., 2012) at default parameter settings (http://cab.spbu.ru/files/release3.11.1/manual.html#correctoropt). The assembled ancient *R. felis* genome was used for average nucleotide identity and single nucleotide variant analysis using the FastANI (Jain et al., 2018) and Snippy (https://github.com/tseemann/snippy) software programmes, respectively. The pan-genome analysis and core genes were identified using 126 Rickettsia species and the BBayA *R. felis* genomes and GET_HOMOLOGUES package (Contreras-Moreira and Vinuesa, 2013) using default parameter settings (https://github.com/eead-csic-compbio/get_homologues). The genome comparison and coverage plots were visualised using the Circos (Krzywinski et al., 2009) package (http://circos.ca/).

### Phylogenetic analysis of BBayA *R. felis*

A concatenated codon alignment was produced from 138 protein sequences using MAFFT v7.464 (Katoh and Standley, 2013) at default parameter settings (https://mafft.cbrc.jp/alignment/software/). The codon alignments were processed using jModelTest (http://evomics.org/learning/phylogenetics/jmodeltest/) to carry out statistical selection of best-fit models of nucleotide substitutions and to select the best model for the phylogenetic analysis. The best model parameters (GTRGI) were used for the construction of maximum likelihood phylogenetic tree using the MEGA X (Kumar et al., 2018) software.

### Molecular clock and divergence analysis of BBayA *R. felis*

Molecular divergence analysis was performed by using the codon alignment in BEAST v2.5.0 (Bouckaert et al., 2019). A coalescent constant prior and strict molecular clock was used for the Markov chain Monte Carlo (MCMC) chain analysis. Five different runs of 100 million MCMC were performed and sampled every 5,000 runs. The independent MCMC runs were combined for the better posterior effective sample size and tree. The starting time for the BBayA *R. felis* was set as 2,000 years BP and all other species were assumed as ‘0’ years. The maximum likelihood tree was supplied as an initial tree for the Bayesian MCMC analysis. The coalescent constant prior and strict clock did not change the topology of the initial tree in the final output. A Burnin tree was produced after discarding the first 10% of trees generated. The final resulting tree was analysed in the iTol tree visualization tool (Letunic and Bork, 2007).

## Supplementary Information (SI 1 - SI 9)

### SI 1 Skeletal provenience of the boy from Ballito Bay

Ballito Boy (BBayA) was recovered from an archaeological context along the KwaZulu-Natal Province coastline and AMS radiocarbon-dated to 1,980 ± 20 cal. BP, *i.e*., *c*. 2,000 years ago (Schlebusch et al., 2017). The remains were excavated by Schoute-Vanneck and Walsh during the 1960s, first curated at the Durban Museum, and then transferred to the KwaZulu-Natal Museum where it is now curated (accession no. 2009/007). The site from which it was retrieved is said to have been a mound formed by a shell-midden overlooking the beach, about 46 m from the high-water mark. The skeletal material cannot be directly associated with archaeological material from the site as clear stratigraphic context is unknown. Admixture analyses indicate that BBayA cluster with modern Southern San populations (Schlebusch et al., 2017). On account of the high genome coverage (∼13-fold) of BBayA, Schlebusch et al. (2017) recalculated the genetic time depth for *Homo sapiens* to between 350 kya and 260 kya. This revised split-estimate coincides with the fossil material from Morocco, dated to *c*. 300 kya (Hublin et al., 2017) and which is viewed as anatomically-transitional between archaic and modern *H. sapiens* (Lombard et al., 2018).

## SI 2 Immune adaptations of African hunter-gatherers

Of the ∼2,100 species of microbes that interact directly with humans (Wardeh et al., 2015), at least 1,415 species are known to be pathogenic, including various bacteria, viruses, fungi, protozoa and helminths (Taylor et al., 2001; Woolhouse and Gowtage-Sequeria, 2005). Approximately 65% of these are zoonotic (Lloyd-Smith et al., 2009) and ∼8% are suspected to cause emerging infectious diseases (Dutour, 2013). At least 20 of these pathogens have certain to probable African origin, including hepatitis B, measles, cholera, dengue fever, *P. falciparum* malaria and leishmaniasis, plague and smallpox (Houldcroft and Underdown, 2016; Wolfe et al., 2007). Despite the fact pathogens have long exerted a significant influence on hominin longevity (Rifkin et al., 2017) and human genetic diversity (Pittman et al., 2016), and given that diseases continue to shape our history (Andam et al., 2016), their influence on the biological and socio-cultural evolution of our species, in Africa, is routinely overlooked.

Persistent exposure to pathogens exerted selective pressure on human health (Owers et al., 2017), immune responses (Nédélec et al., 2016), cognitive development (Kessler et al., 2017) and social behaviour (Thornhill and Fincher, 2014). The bio-geographic distribution of *Plasmodium falciparum* (Tanabe et al., 2010) and *Helicobacter pylori* (Linz et al., 2007) exhibits declining genetic diversity, with increasing distance from Africa, with ‘Out of Africa’ estimates of about 58 kyr and 80 kyr ago, respectively. Indeed, and given that the *H. pylori* association with humans is at least 100,000 years old (Moodley et al., 2012), the current population structure of *H. pylori* may be regarded as mirroring past human expansions and migrations. In addition to *Plasmodium falciparum* (Tanabe et al., 2010), roughly 250 *Plasmodium* species, including *P. vivax, P. malariae, P. falciparum* and *P. ovale* are highly anthropophilic (Ollomo et al., 2009). Mitochondrial mtDNA analyses confirm that early forms of *P. falciparum* were present by at least 100 kya (Kwiatkowski, 2005; Silva et al., 2011). Some of the first examples of natural selection acting on the human genome involve genetic mutations that confer resistance to malaria. The Duffy negativity locus evolved some 100 ka (Ferwerda et al., 2007) to ∼ 60 kya (McManus et al., 2017) and confers resistance against *P. vivax* malaria to many sub-Saharan Africans (Howes et al., 2011). That these and several other malaria-resistant alleles evolved independently (Ko et al., 2011) suggests that malaria exerted a significant degree of selective pressure in prehistory.

In addition to the fact that it appears that persistent exposure to pathogens exerted selective pressure on human immune-related genes (Nédélec et al., 2016; Owers et al., 2017), the antiquity of genetic disease prevention mechanisms, such as the origin of immune-regulating Sia-recognising Ig-like lectin (SIGLEC) genes before 70 kya (Wang et al., 2012), confirms that pathogens played an essential role in human evolution in Africa. More recently, Lopez et al. (2019) has detected strong polygenic adaptation signals for functions related to mast-cell responses to allergens and microbes, and host interactions with viruses also support a history of pathogen-driven selection in the rainforest. In the case of BBayA, the incidence of genomic variants relating to pathogen exposure (Schlebusch et al., 2017) is of particular interest. The FY*A allele, which has a protective effect against malaria, was identified in BBayA, which also carries the ATP2B4 gene variant, another polymorphism which protect against childhood malaria and which appears to have emerged ∼ 60 kya (McManus et al., 2017). BBayA does not carry the Duffy null allele, which has a protective effect against *P. vivax* associated malaria. Similarly, the APOL1 gene variant, which confers resistance to African sleeping sickness, is also absent in BBayA.

## SI 3 *Rickettsia felis* strain LSU-Lb

*Rickettsia felis* str. LSU-Lb is an obligate mutualist of the parthenogenic booklouse *Liposcelis bostrychophila* (Insecta: Psocoptera), an insect only recently recognized as a host for *R. felis* (Thepparit et al., 2011). *Rickettsia felis* str. LSU-Lb was first isolated in 2010, in Los Angeles County, California, USA. Phylogenomic analysis suggests that *R. felis* str. LSU-Lb diverged from the flea-associated strains. It is suggested that the shared microhabitat between fleas (*e.g*., cat fleas, *Ctenocephalides felis*) and *L. bostrychophila* and the phoretic relationship of *R. felis*-infected *L. bostrychophila* with vertebrate hosts facilitates the horizontal transmission of *R. felis* from fleas to *L. bostrychophila*.

## SI 4 *Toxoplasma gondii* TgCatPRC2

*Toxoplasma gondii* was first isolated in the early 1900s from a North African rodent, the common gundi (*Ctenodactylus gundi*), from which the species name was derived. Subsequent surveys have found *T. gondii* to be highly prevalent among many species of mammals and birds (Su et al., 2012). In contrast to Europe and the Americas, the genetic diversity and population structure of *T. gondii* from Africa, where limited data are available, remains controversial (Mercier et al., 2010). Two recent genotyping studies based on African *T. gondii* strains have suggested that, like in Europe and in the USA, the same three main lineages predominate in Africa, with one strain considered to be a recombinant between Type II and III strains. Non-classical genotypes of *T. gondii*, called *Africa 1, Africa 2* and *Africa 3* have been isolated from immunocompromised patients with toxoplasmosis acquired in Western and Central Africa. Because these genotypes were also recovered in patients from different African countries, they were proposed as common clonal lineages in Africa.

It is clear that the population structure of *T. gondii* in Africa is far from being resolved. It appears that the *Africa 1* strains and the single genotype clusters with Type I and Type II reference strains, respectively (Galal et al., 2019). *Africa 1* also clusters with GPHT, DPHT, TgCkBr59, and TgCkBr40 strains. *Africa 3* clusters with TgCkBr93, and Type III and III-like strains with Type III reference strains. ENVL-2002-MAC strain was closed to one of the Type III-like strains (GAB4-2007-GAL-DOM1). The *Africa 2* strain (CCH002-2004-NIA) is different to all the other strains. Whether these additional haplogroups for Africa represent minor variations of Type I, II, and III, or recombinant strains of these three lineages, remains to be determined. Strains from China have been found to comprise haplogroups distinct from those previously described (Khan et al., 2011). Strains TgCtPRC2 and TgCtPRC6 were identical to each other and represent a common genotype in China. The *T. gondii* TgCatPRC2 strain, obtained from a cat in Guangzhou, Guangdong Province, China, in 2007, was sequenced as the reference strain of *T. gondii* haplogroup 13 at the J. Craig Venter Institute (https://www.ncbi.nlm.nih.gov/bioproject/167493).

## SI 5 Do bacteria have lower aDNA damage patterns than eukaryotes?

Following DNA extraction, the sequencing output, *i.e*., ‘read-counts’, is dependent on the sequencing depth of each sequencing run and the presence of sufficient un-damaged DNA strands to detect during sequencing, the latter factor which is, in turn, dependent on the morphology (*i.e*., the cell wall structure, spore formation, the presence of mycolic acids and guanine-cytosine (GC) content) of different types of pathogenic microbes (Mann et al., 2018). As per Donoghue et al. (2017), mycobacterial aDNA is generally more resistant to degradation compared to mammalian host aDNA, due to the protective presence of the bacterial cell wall and the higher proportion of guanidine and cytosine in the DNA. However, Mann et al. (2018) found that fragmentation patterns within dental calculus are associated with the genomic source of the DNA (human *vs*. microbial) but not with cellular structure (*e.g*., microbial cell wall type or presence of a surface-layer). Accordingly, it appears that short DNA fragments from taxa with lower GC content genomes should be expected to be more susceptible to loss through denaturation because their melting point for a given fragment length will be lower, and this may contribute significantly to taxonomic misalignments and misidentifications.

Consistent with this hypothesis, Mann et al. (2018) found that high GC-content genera had slightly shorter median fragment lengths overall, which accords with a higher retention of short DNA fragments.

## SI 6 The emergence of a MRCA for the southern African *R. felis* group

Our analyses also revealed the emergence of a most recent common ancestor (MRCA) for the southern African *R. felis* group at 5,000 ya (*i.e*., 5,282 ya - 4704 ya), during the microlithic Wilton techno-complex of the southern African Later Stone Age (LSA) (Lombard et al., 2012). The Wilton spanned the period from approximately 8,000 to 4,000 years ago and is characterised by a fully developed microlithic tradition with numerous formal tools, highly standardised backed microliths and small convex scrapers, the widespread use of ostrich eggs and also egg-shell beads, the habitual exploitation of red ochre and the common use of tools made from bone, wood and also shell (Lombard et. al., 2012).The Wilton is also marked by a significant increase (*i.e*., by ∼45%) in archaeological sites (*n* = 34, including Blombosfontein, Boomplaas Cave, Buzz Shelter, Byneskranskop, Cave James, Diamond Shelter, Dikbosch, Elands Bay Cave, Gehle Shelter, Good Hope Shelter, Jakkalsberg, Jubilee Shelter, Kabeljous River Shelter, Kangkara Cave, Kasteelberg, Leliehoek, Maqonqo Shelter, Matjes River, Melkhoutboom Cave, Mgede Shelter, Mzinyashana Shelter, Nelson Bay Cave, Nkupe Shelter, Oakhurst Cave, Rooikrans Shelter, Rose Cottage Cave, Sehonghong, Springbokoog, Steenbokfontein, The Havens Cave, Tloutle Rock Shelter, Tshisiku Shelter, Wilton Large Rock Shelter and Wonderwerk Cave), and, by proxy, human population density. The divergence time for *R. felis* URRWxCal2 and *R. felis* BBayA was estimated at 2,942 ya and that of *R. felis* LSU-Lb and *R. felis* BBayA at *c*. 2,000 ya.

## SI 7 Pathogenicity and clinical symptoms of *Rickettsia felis* infection

*Rickettsia felis*, an insect-borne rickettsial pathogen and the causative agent of typhus-like flea-borne ‘spotted fever’, is an obligate intracellular bacterium in the order Rickettsiales (Angelakis et al., 2016). While cat- and dog-fleas (*Ctenocephalides felis* and *C. canis*) have been cited as the most probable vectors, >40 different haematophagous species of fleas, mosquitoes, ticks and mites have been identified as vectors (Legendre and Macaluso, 2017). As well as the identification of the African great apes (chimpanzees, gorillas, and bonobos) as vertebrate reservoirs responsible for the maintenance of *R. felis* in Africa, it has been proposed that humans are natural *R. felis* reservoirs (Mediannikov et al., 2014), just as they are for certain *Plasmodium* species (Gonçalves et al., 2017).

*R. felis* is therefore capable of infecting multiple hosts and vectors, and co-feeding likely explains the enzootic spread of *R. felis* among variable host- and vector-populations (Angelakis et al., 2016; Brown and Macaluso, 2016). In addition, while rickettsial diseases are widely stated to represent emerging infectious pathogens, the historic influence of *Rickettsia* is well-known. Whereas the first evidence of *R. felis’s* potential as a human pathogen surfaced in 1994 (Angelakis et al., 2016), the first reliable description of typhus-like disease appears in 1489 during the Spanish siege of Baza against the Moors during the War of Granada (1482 to 1492) (Pages et al., 2010). Ancient DNA analysis of human remains and body lice (*Pediculus humanus*) recovered from the graves of soldiers who perished during Napoleon’s 1812 Russian Campaign, confirmed historic accounts of the presence of both trench fever (*Bartonella quintana*) and epidemic typhus (*Rickettsia prowazekii*) during the campaign (Raoult et al., 2006).

The clinical presentation of rickettsial diseases ranges from mild to severe. Without antibiotic treatment, murine or ‘endemic’ typhus, caused by *R. typhi*, exhibits a mortality rate of 4%, and Rocky Mountain spotted fever a mortality rate as high as 30% (Snowden and Bhimji, 2017). Epidemic typhus, caused by *R. prowazekii*, has a mortality rate which varies from 0.7% to 60% for untreated cases. Mortality rates as high as 66% has been reported for disease due to *R. rickettsii* occurring prior to 1920, preceding the discovery of antibiotics (Azad, 2007). The minimal genomic divergence distinguishing *R. felis* from other flea-associated strains suggests that it has the potential to be a human pathogen (Gillespie et al., 2015). The clinical manifestations of *R. felis* infection closely resemble those of flea-borne murine typhus (Blanto and Walker, 2016) which entails the abrupt onset of fever with accompanying headache, chills, myalgia, malaise and cutaneous maculopapular rashes (Angelakis et al., 2016; Legendre and Macaluso, 2017).

The similarity of typhus-like flea-borne rickettsioses symptoms to *R. typhi*, as well as the lack of specific diagnostics, has potentially resulted in the under-diagnosis of *R. felis* in many human cases (Legendre and Macaluso, 2017). In sub-Saharan Africa, *R. felis* is described as a common (∼15 %) cause of illness among patients with ‘fever of unknown origin’, particularly in malaria-endemic regions (Brown and Macaluso, 2016). In some regions, the incidence of human *R. felis* infections far-exceeds that of malaria. Diagnosis is problematic because symptoms are common to other infectious diseases, including mosquito-borne dengue fever (Flavivirus) and malaria (e.g., *P. falciparum*) and brucellosis (*B. melitensis*). *R. felis* has furthermore been detected in the blood and cerebrospinal fluid of those with an alternative and more compelling diagnosis, including malaria, cryptococcal meningitis and scrub typhus (Blanton and Walker, 2017). The clinical presentation of rickettsial diseases can vary from mild to very severe, with the case-fatality rate for highly virulent rickettsiae ranging from 2% to 30% (Azad, 2007). Human disease case fatality rates (CFRs), the proportion of patients that reportedly died as a result of infection, of 19% have been reported for untreated *R. felis* infections (Oliveira et al., 2002).

## SI 7 Pathogenicity and clinical symptoms of *Toxoplasma gondii* infection

*Toxoplasma gondii* is perhaps the best known of the tissue cyst-forming coccidians, a group of organisms that have heteroxenous life cycles that alternate between sexual replication in the intestinal epithelium of their definitive hosts, and asexual replication that occurs in tissues of intermediate hosts.

*T. gondii* is unusual in being a generalist among this group, with members of the cat family (Felidae) serving as the definitive hosts, while seemingly all homoeo-thermic vertebrates can serve as intermediate hosts (Behnke et al., 2016). *T. gondii* it is estimated to chronically infect one-third of the global human population, causing ocular toxoplasmosis, encephalitis as well as birth defects following vertical transmission to developing foetuses (Shwab et al., 2018). Although latent toxoplasmosis is generally assumed to be asymptomatic in immunocompetent individuals, it can induce several behavioural manipulations in infected humans. Dubbed the human ‘brain parasite’, *T. gondii* displays prominent tropism for the brain tissue, where it forms intracellular cysts within the neurons and glial cells, establishing a chronic infection. Chronic *T. gondii* infection in the human population has been correlated with a diverse range of human diseases, including Alzheimer’s and Huntington’s. The association between *T. gondii* infection and mental health disorders such as depression, psychosis, self-directed violence, bipolar disorder and schizophrenia have been widely studied (Tedford and McConkey, 2017).

Serological studies demonstrate its presence in virtually every country, with seroprevalence exceeding 60% in parts of South America, Africa, and South-East Asia (Pappas et al., 2009). Various factors underlie the severity and clinical heterogeneity of toxoplasmosis, such as the mode of contamination, the inoculum dose, repeated *T. gondii* infections, age, ethnic variation, immune status, occurrence of co-infections and genotypes (Galal et al., 2019). Variability in human infection and mortality is largely ascribed to the causative *T. gondii* strain type. Several lineages present greater clinical severity in otherwise healthy humans. For example, type 4 and related type 8 strains are associated with severe ocular infections, and types 5 and 10 can cause serious disease in otherwise healthy adults (Behnke et al., 2016). Sampling from Africa is sporadic, but at least one haplotype commonly found in Africa (*i.e*., type 14) is related to type 6 strains found in both Europe and South America, suggesting a global distribution for this clade.

Congenital toxoplasmosis can result in some of the most serious consequences with a wide range of clinical manifestations, including spontaneous abortions and stillbirths, or in live infants, hydrocephalus, microcephaly, retinochoroiditis and cerebral calcifications (Hammond-Aryee et al. 2014). Clinical manifestation of toxoplasmosis because of immunosuppression, typically due to reactivation of a chronic infection or acquisition of a new infection, manifests as toxoplasmic encephalitis (TE), also known as central nervous system toxoplasmosis (Hammond-Aryee et al. 2014). Disease progression leads to severe manifestations such as confusion, lethargy, mental state changes, seizures, and coma, and the outcome is fatal. Given the extraordinary implications of *T. gondii* infection for bizarre human behaviours, it is surprising that the co-evolutionary history of the parasite and its intermediate human hosts remains shrouded in mystery. Whereas case fatality rates (*i.e*., the proportion of patients that reportedly died as a result of infection) of 29% have been reported for infections amongst immunocompromised (HIV-AIDS) patients (Luma et al., 2013), misdiagnosis and comorbidities likely result in low case fatality estimations in otherwise healthy individuals. CFRs of ∼1.0% have been reported for a Tanzanian hospital-based (*i.e*., admitted) survey (Mboera et al., 2019), and a CFR of 25.5% was reported by a study in the USA during the period from 2000 to 2010 (Cummings et al., 2014).

## SI 8 Pathogenicity and clinical symptoms of *Anaplasma, Babesia, Bordetella, Brucella, Leishmania, Plasmodium* and *Trypanosoma* infection

At the genus level, we identified aDNA reads mapping to *Anaplasma, Babesia, Bordetella, Brucella, Leishmania, Plasmodium* and *Trypanosoma* (Table S1). The authentication of aDNA sequence reads ascribed to these taxa was achieved by library-independent verification using mapDamage (Jónsson et al., 2013) and analyses of the read-length distribution (bp) (Fig. S1).

*Anaplasma* is a bacterium and the causative agent of anaplasmosis. It is spread to humans by tick bites primarily from the *Ixodes* genus (see https://www.cdc.gov/anaplasmosis/index.html). Early signs and symptoms occurring 1 to 5 days after initial infection are usually mild or moderate and may include fever, chills, severe headache, muscle aches nausea, vomiting, diarrhoea and loss of appetite. More severe symptoms, and if treatment is delayed or if there are other medical conditions present, can result in severe illness. The signs and symptoms of severe (late stage) illness include respiratory failure, bleeding problems, organ failure and death. While as many as 3.0% of infected individuals may develop life threatening complications, CFRs of ∼1.0% have been reported hospitalised anaplasmosis patients (Bakken and Dumler, 2014).

Because *Babesia* parasites infect and destroy red blood cells, babesiosis causes hemolytic anaemia (https://www.cdc.gov/parasites/babesiosis/disease.html). This type of anaemia can lead to jaundice (yellowing of the skin) and dark urine. Further complications include low and unstable blood pressure, severe hemolytic anaemia (haemolysis), a very low platelet count (thrombocytopenia), disseminated intravascular coagulation (consumptive coagulopathy), which can lead to blood clots and bleeding, the malfunction of vital organs (such as the kidneys, lungs, and liver) and death. CFRs of 9.0% have been reported for infected hospitalised individuals (Vannier et al., 2008).

*Bordetella*, of which *Bordetella pertussis* is the causative agent of whooping cough, cause serious illness in children, teens and adults (https://www.cdc.gov/pertussis/about/signs-symptoms.html). Symptoms develop within 10 days after exposure. After 1 to 2 weeks, and as the disease progresses, the traditional symptoms of pertussis may appear and include paroxysms (fits) of many, rapid coughs followed by a high-pitched ‘whoop’ sound, vomiting (throwing up) during or after coughing fits and exhaustion after coughing fits. There are several antibiotics available to treat pertussis. CFRs of 4.0% have been reported for infections in developing countries (Gabutti and Rota, 2012).

Brucellosis is a severely debilitating bacterial disease caused by members of the genus *Brucella* (https://www.cdc.gov/brucellosis/symptoms/index.html). Humans are infected via contact with infected animals or animal products. Some signs and symptoms may persist for longer periods of time, including recurrent fevers, arthritis, swelling of the heart (endocarditis), neurologic symptoms (in up to 5% of all cases), chronic fatigue, depression and also swelling of the liver and spleen.

Depending on the timing of treatment and severity of illness, recovery may take a few weeks to several months. Although death from brucellosis is rare, occurring in no more than 2% of all cases, CFRs as high as 6.5% have been reported (Dahouk et al., 2007).

Leishmaniasis is a parasitic disease caused by infection with *Leishmania* parasites spread by the bite of phlebotomine sand flies (https://www.cdc.gov/parasites/leishmaniasis/). There are several forms of leishmaniasis, with the most common being cutaneous leishmaniasis, which causes skin sores, and visceral leishmaniasis, which affects internal organs, usually the spleen, liver and bone marrow. If not treated, severe cases of visceral leishmaniasis are fatal. Globally, CFRs of 10.0% have been reported for documented (*i.e*., hospitalised) infections (Alvar et al., 2012).

Approximately 156 members (*i.e*., species) of the genus *Plasmodium* are the causative agents of malaria (https://www.cdc.gov/dpdx/malaria/). The clinical presentation of malaria infection can vary substantially depending on the infecting species, the level of parasitemia and the immune status of the individual. Untreated malaria progress to severe forms that may be rapidly (*i.e*., within 24 hours) fatal. Infections caused by *P. falciparum* are the most likely to progress to severe, potentially fatal forms with central nervous system involvement (cerebral malaria), acute renal failure, severe anaemia or acute respiratory distress syndrome. Other species can also have severe manifestations, such as *P. vivax* malaria, which include splenomegaly (including splenic rupture), and those of *P. malariae* which include nephrotic syndrome. CFRs of 15.0% (Camponovo et al., 2017) to 35.0% (Kapesa et al., 2018) (*c*. 25%) have been reported for individuals hospitalised in malaria-endemic African regions.

African trypanosomes are protozoan haemoflagellates of the genus *Trypanosoma*. Two subspecies cause disease in humans, including *T. b. gambiense*, causing chronic African trypanosomiasis or ‘West African sleeping sickness’ and *T. b. rhodesiense*, causing acute African trypanosomiasis or ‘East African sleeping sickness’ (https://www.cdc.gov/dpdx/trypanosomiasisafrican/). The first stage of the disease (*i.e*., the haemolymphatic stage) involves non-specific signs and symptoms such as intermittent fever, pruritus and lymphadenopathy. During the second stage of the disease, the meningoencephalitic stage, invasion of the central nervous system causes a variety of neuropsychiatric manifestations, including sleep disorders. Severe cardiac involvement with electrocardiogram abnormalities consistent with perimyocarditis is also observed. CFRs of 17.2% have been reported for hospitalised with advanced infections (Kazumba et al., 2018).

## SI 9 The most plausible range of microbes implicated in the death of the boy from Ballito Bay

In the case of the child from Ballito Bay, the data presented is certainly indicative of the extreme severity of his co-infective clinical condition. One would be inclined to infer that infection by one pathogen would, almost inevitably, facilitate increasing susceptibility to subsequent infection by other pathogenic taxa. However, determining the consequences of co-infection is marred by problems as there are many confounding factors to consider (McArdle et al., 2018), particularly as co-infections may be insignificant, detrimental or even beneficial, and because these outcomes depend on various levels of interactions between pathogenic taxa and the host immune system.

It is, consequently, difficult to determine the exact impact of the identified ancient pathogenic taxa on the morbidity and mortality of the boy from Ballito Bay. First, the total number of unique ancient (*i.e*., authenticated) aDNA sequence reads cannot be viewed as representative of the severity of infection of a particular pathogen. It has been shown that different DNA pre-processing and extraction methods result in notable differences in the microbial profiles generated (Eriksen et al., 2020). As indicated above, following DNA extraction, the sequencing output (‘read-counts’) is dependent on the sequencing depth of each sequencing run and the presence of sufficient un-damaged DNA strands to detect during sequencing. The latter factor is dependent on variable factors, including the cell wall structure, spore formation ability, the presence of mycolic acids and also the guanine-cytosine (GC) content of different types of pathogenic microbes (Mann et al., 2018). In addition, the relationship between ‘pathogen load’ and severity is difficult to determine as pathogens can be distributed, non-uniformly, throughout multiple different cell, tissue or organ compartments of the body (Cunnington, 2015). Furthermore, as the DNA analysed in this study was extracted from bone (petrous) and tooth (upper premolar) samples, the fact that not all pathogenic microbial taxonomic categories might be recoverable from either human skeletal or dental remains (Margaryan et al., 2018) suggest that there is a possibility that some taxa might be underrepresented. In this regard, and as opposed to the results reported by Margaryan et al. (2018), our study confirms that the DNA of various ancient pathogenic microbial taxa can indeed be recovered from human petrous bone samples.

Second, we depend on CFRs reported for contemporary hospital settings, in which patients were subjected to treatment, and not on the CFRs of these pathogens in prehistoric contexts during which no modern medicinal treatments (*i.e*., diagnostic equipment and antibiotics) would have been available. Third, and although ancient human co-morbidities can be inferred from pathogenic aDNA (Houldcroft et al., 2019) it is highly probable that ancient genomic traces of viral taxa, particularly that derived from single-stranded RNA viruses such as Yellow fever (of the family Flaviviridae) and Hepatitis A (Picornaviridae) implicated in the co-infection of the child, was simply not detected.

Fourth, many fundamental patterns of pathogen co-infection (*i.e*., multi-species infections) remain undescribed, including the relative frequency of co-infection by various pathogens and the differences between single-species infections and co-infection. Although current clinical research do indicate marked differences in the morbidity of co-infected patients, as opposed to those with single infections (with co-infection having seriously adverse health implications), the effects of co-infection are generally interpreted relative to the clinical conditions observed during infections of single pathogen species (Griffiths et al., 2011). A fifth confounding factor involves the fact that ‘within-host’ ecological competition between infective pathogens poses another problematic caveat, especially given the observation that co-infecting pathogens do not always interact or compete with each other (Hamelin et al., 2019). Sixth, and in addition to the combined influence of the CFRs reported for the detected ancient pathogenic taxa, it must be noted that a diagnosis of anaemia, unaccompanied by other infections, has been reported to have a post-hospitalisation mortality rate of ∼20.0% for sub-Saharan African children (Chami et al., 2019). Finally, it is evident that, in order to be accurate in its species assignment, the reference database used by Kraken2 must be highly representative of the described species and strains. The more exhaustive the database, the better the assignment accuracy will be, and the lack of African reference genomes for various taxa, is therefore highly problematic.

## Supplementary Figures

**Figure S1.**
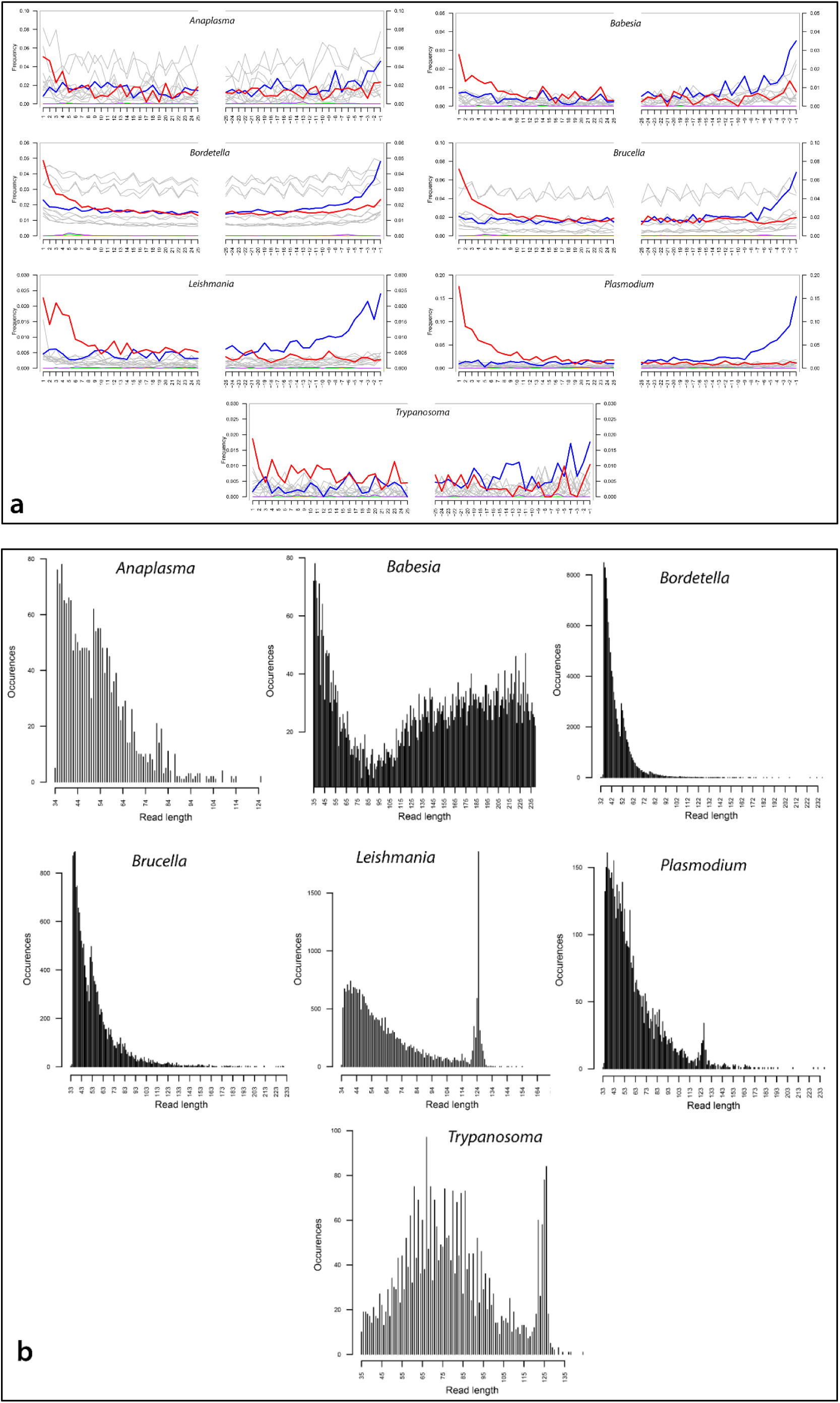
**a**) DNA damage pattern analysis for pathogenic members of the BBayA parasitic genera *Anaplasma, Babesia, Bordetella, Brucella, Leishmania, Plasmodium* and *Trypanosoma* using mapDamage. G-to-A and C-to-T misincorporations are plotted in blue and red, respectively. Grey lines indicate all possible misincorporations. **b**) DNA fragment read-length distributions for *Anaplasma* (with a mean read-length of 70.17 bp), *Babesia* (187.57 bp), *Bordetella* (56.29 bp), *Brucella* (67.83 bp), *Leishmania* (67.88 bp), *Plasmodium* (61.77 bp) and *Trypanosoma* (77.79 bp) reads.

**Figure S2.**
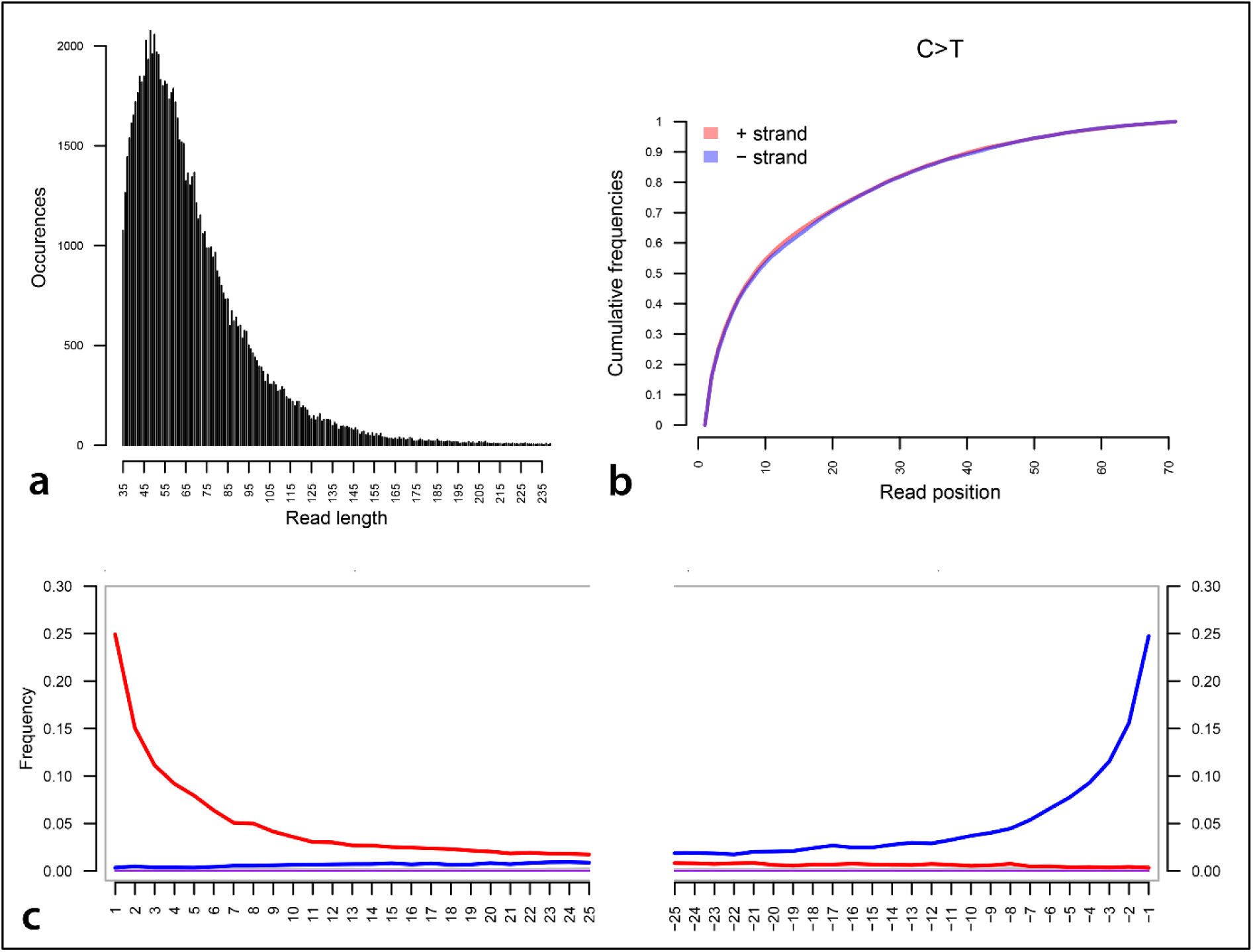
Damage pattern and read-length distribution analysis of the human host’s (BBayA) DNA exhibit a similar DNA damage profile and short (*i.e*., damaged) read-length distribution to that of the *R. felis* and *T. gondii* DNA sequence reads analysed. **a**) DNA fragment read-length distributions of the BBayA host reads, **b**) C-T read strand positions and **c**) G-to-A and C-to-T misincorporations are plotted in blue and red, respectively, and the grey lines indicate all possible misincorporations.

**Figure S3.**
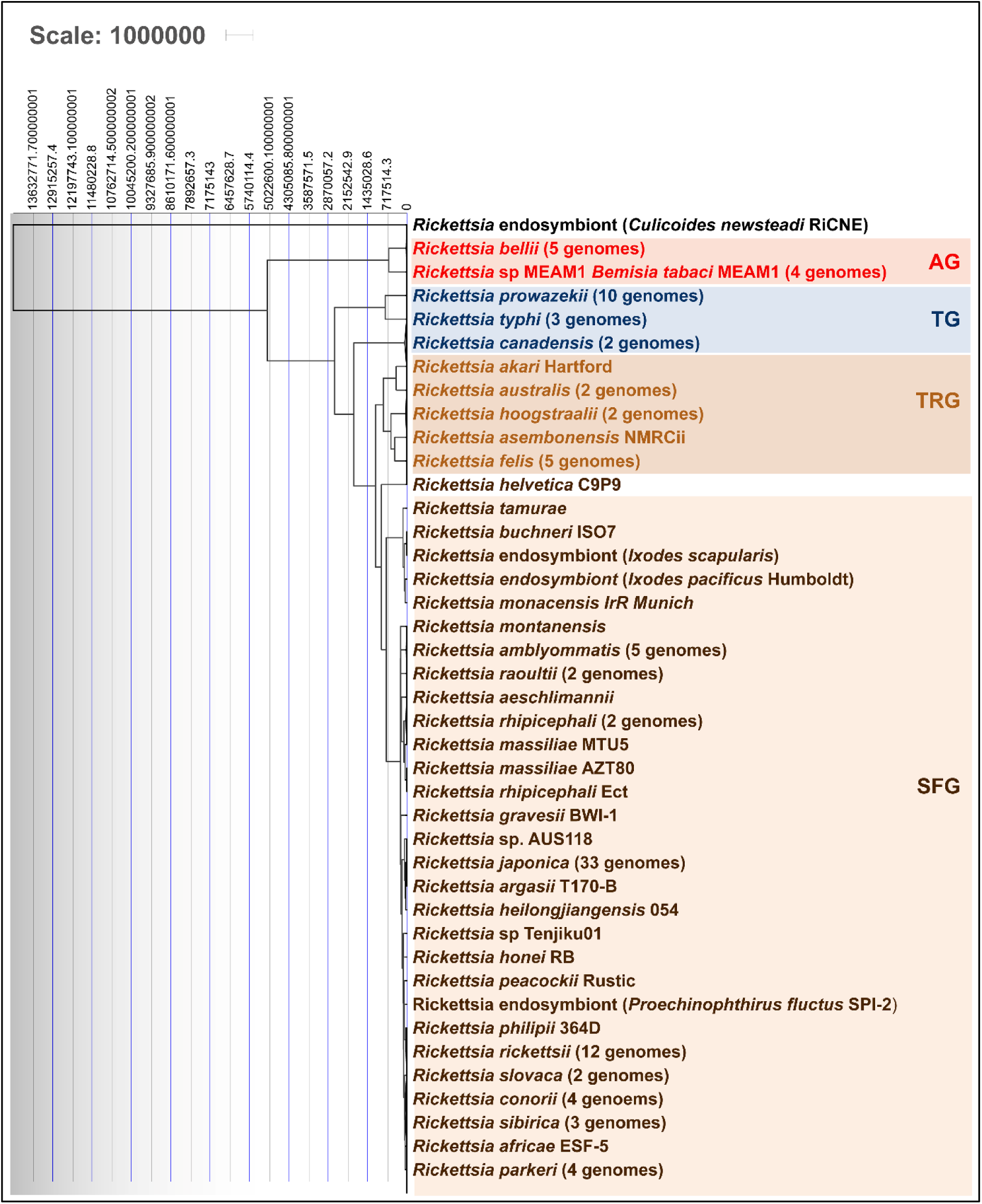
Molecular clock and divergence time analysis of the ancient BbayA *R. felis* genome was performed using all (*i.e*., 126) currently available NCBI reference genomes. The MCMC algorithm was applied on the codon alignment of 138 core genes using a strict clock, a coalescent constant and the GTRGI substitution model in Beast2.

**Figure S4.**
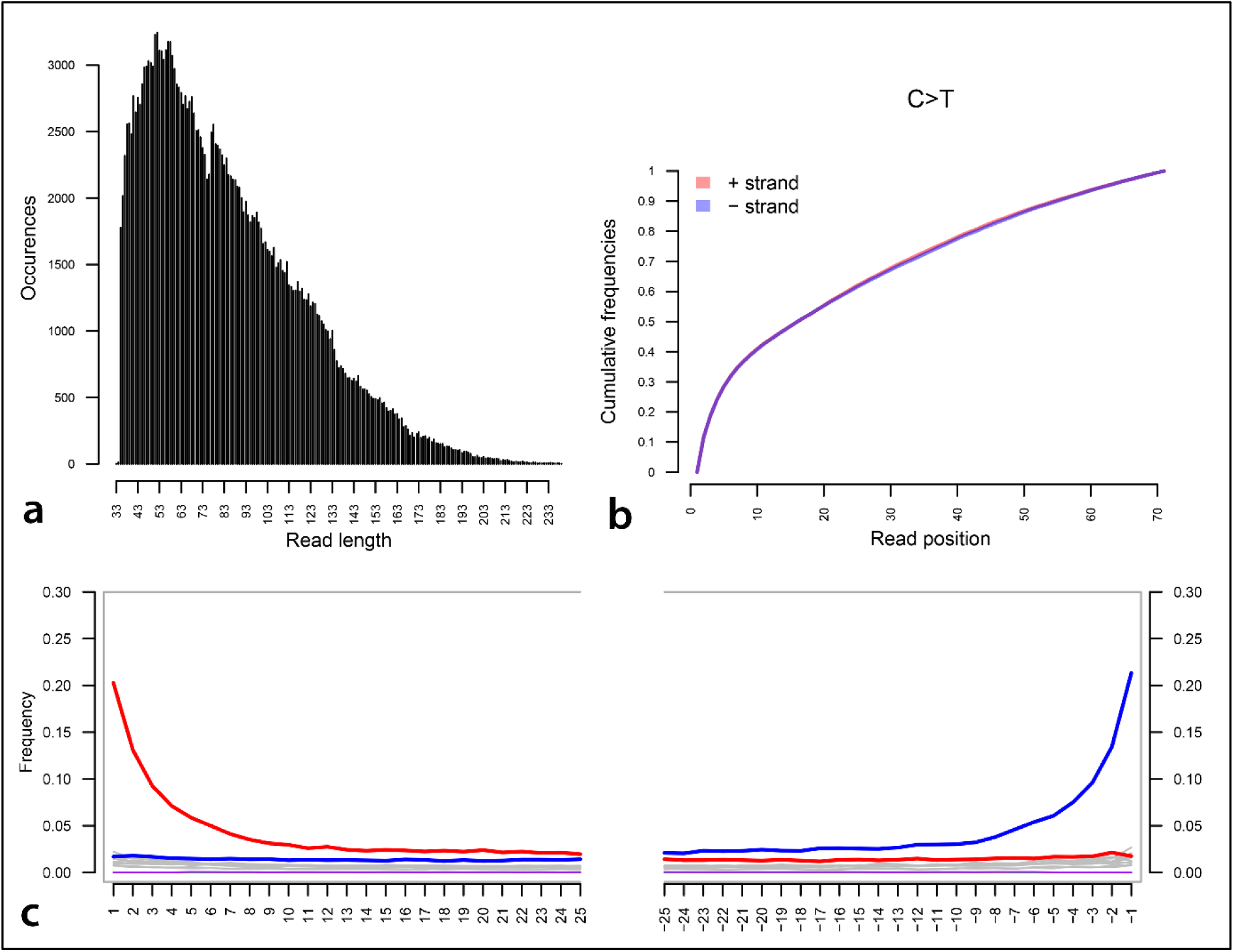
**a**) DNA damage pattern analysis for the ancient BbayA *T. gondii* reads. **a**) DNA fragment read-length distributions of the BBayA host reads, **b**) C-T read strand positions and **c**) G-to-A and C-to-T misincorporations are plotted in blue and red, respectively, and the grey lines indicate all possible misincorporations.

## Supplementary Tables

**Table S1**. Additional pathogenic taxa (*i.e*., *Anaplasma, Babesia, Bordetella, Brucella, Leishmania, Plasmodium* and *Trypanosoma*) detected in the BBayA child, indicating the percentage of total metagenomic reads, the percentage of read duplicates, total number of unique reads and the mean read-length (bp). Competitive alignment of short reads was performed to determine the those mapping to pathogenic taxa. Average genome coverage was calculated using the unique read alignments and standard genome sizes.

**Table S2**. Mapping of the BBayA aDNA sequence dataset was performed on a competitive basis against bacterial and parasitic genomes, and a complete human genome. NCBI reference assembly genomes are indicated for all the authenticated taxa detected in the BBayA metagenomic dataset.

**Table S3**. The 126 NCBI reference genomes initially used to identify the closest genomic homologues to the ancient BBayA *R. felis* strain.

**Table S4**. The NCBI reference genomes used for phylogenetic analyses and comparison of the BBayA *R. felis* to *R. felis* LSU-Lb, *R. felis* URRWxCal2, *R. typhi, R. prowazekii* and *R. africae*.

**Table S5**. Reported disease case fatality rates (CFRs) for pathogenic members of the microbial genera identified in the BBayA child illustrating the severity of his co-infective clinical condition.

